# Bacterial-like nonribosomal peptide synthetases produce cyclopeptides in the zygomycetous fungus *Mortierella alpina*

**DOI:** 10.1101/2020.08.21.262279

**Authors:** Jacob M. Wurlitzer, Aleksa Stanišić, Ina Wasmuth, Sandra Jungmann, Dagmar Fischer, Hajo Kries, Markus Gressler

## Abstract

Fungi are traditionally considered as reservoir of biologically active natural products. However, an active secondary metabolism has long not been attributed to early diverging fungi such as *Mortierella spec*. Here, we report on the biosynthesis of two series of cyclic pentapeptides, the malpicyclins and malpibaldins, as products of *Mortierella alpina* ATCC32222. The molecular structures of malpicyclins were elucidated by HR-MS/MS, Marfey’s method, and 1D and 2D NMR spectroscopy. In addition, malpibaldin biosynthesis was confirmed by HR-MS. Genome mining and comparative qRT-PCR expression analysis pointed at two pentamodular nonribosomal peptide synthetases (NRPS), malpicyclin synthetase MpcA and malpibaldin synthetase MpbA, as candidate biosynthetic enzymes. Heterologous production of the respective adenylation domains and substrate specificity assays proved promiscuous substrate selection and confirmed their respective biosynthetic roles. In stark contrast to known fungal NRPSs, MpbA and MpcA contain bacterial-like dual epimerase/condensation domains allowing the racemization of enzyme-tethered l-amino acids and the subsequent incorporation of d-amino acids into the metabolites. Phylogenetic analyses of both NRPS genes indicate a bacterial origin and a horizontal gene transfer into the fungal genome. This is the first report of nonribosomal peptide biosynthesis in basal fungi which highlights this paraphylum as novel and underrated resource of natural products.

**IMPORTANCE:** Fungal natural compounds are industrially produced with application in antibiotic treatment, cancer medications and crop plant protection. Traditionally, higher fungi have been intensively investigated concerning their metabolic potential, but re-identification of already known compounds is frequently observed. Hence, alternative strategies to acquire novel bioactive molecules are required. We present the genus *Mortierella* as representative of the early diverging fungi as an underestimated resource of natural products. *Mortierella alpina* produces two families of cyclopeptides, denoted malpicyclins and malpibaldins, respectively, via two pentamodular nonribosomal peptide synthetases (NRPSs). These enzymes are much closer related to bacterial than to other fungal NRPSs, suggesting a bacterial origin of these NRPS genes in *Mortierella*. Both enzymes are the first biochemically characterized natural product biosynthesis enzymes of basal fungi. Hence, this report establishes early diverging fungi as prolific natural compound producers and sheds light on the origin of their biosynthetic capacity.

## Introduction

The increasing number of human pathogenic microbes resistant against well-established antibiotics requires an unnbiased search of novel producers of bioactive natural products. Traditionally, bacteria (e.g. *Actinomyces* or *Streptomyces* spp.) and filamentous higher fungi (e.g. *Aspergillus*, *Penicillium*, *Fusarium*) are well-investigated resources of natural products such as nonribosomal peptides (NRP), polyketides or terpenoids (1–4). In contrast, early diverging fungi - formerly combined in the paraphylum “zygomycetes” (5, 6) - have long been thought to lack secondary metabolites (7). Still, apart from the C-18-terpenoid trisporic acid isolated from *Blakeslea trispora* and *Mucor mucedo* as a signaling compound during mating (8–10), secondary metabolites are rarely observed in zygomycetes. *Mortierella alpina*, a species of the subdivision *Mucoromycotina*, is a strain that is “generally regarded as safe” (GRAS) and serves as an industrial producer of polyunsaturated fatty acids such as arachidonic acid or linoleic acid (11–13).

Although lipid extracts of *M. alpina* are useful as immunomodulating leukotriene precursor molecules in pharmaceutical applications or as nutritional supplements in baby food (14), the secondary metabolome of the fungus has never been investigated in detail. Case reports on *M. alpina* species indicate the capacity to produce some oligopeptides such as calpinactam (15) and Ro 09-1679 (16) harboring antimycobacterial and thrombin-inhibiting bioactivity, respectively (Figure 1A). The metabolic potential of *M. alpina* was expanded by the recent discovery of a family of linear, acetylated hexapeptides, malpinins A-D, that have surface tension lowering properties (Figure 1A) (17, 18). Furthermore, the fungus produces the highly hydrophobic cyclopentapeptides, malpibaldins (compounds **1-3**) (17). Cyclopentapeptides have been isolated from various biological sources (ascomycetes, algae and bacteria) and harbor diverse pharmaceutically relevant antiviral, antibiotic, apoptotic or anti-angiogenic properties (Figure 1B) (19–25). Of interest, malpibaldins from *M. alpina* are structurally similar to luminmide B, a d-amino acid containing cyclic pentapeptide from the insect pathogenic γ-proteobacterium *Photorhabdus luminescens* (24). Biosynthetically, luminmide B is produced by a pentamodular NRP synthetase (NRPS), Plu3263, by sequential condensation of five aliphatic l-amino acids activated by specific adenylation (A) domains and d-amino acids are introduced by the activity of dual epimerization/condensation (E/C) domains (24).

**Figure 1.**
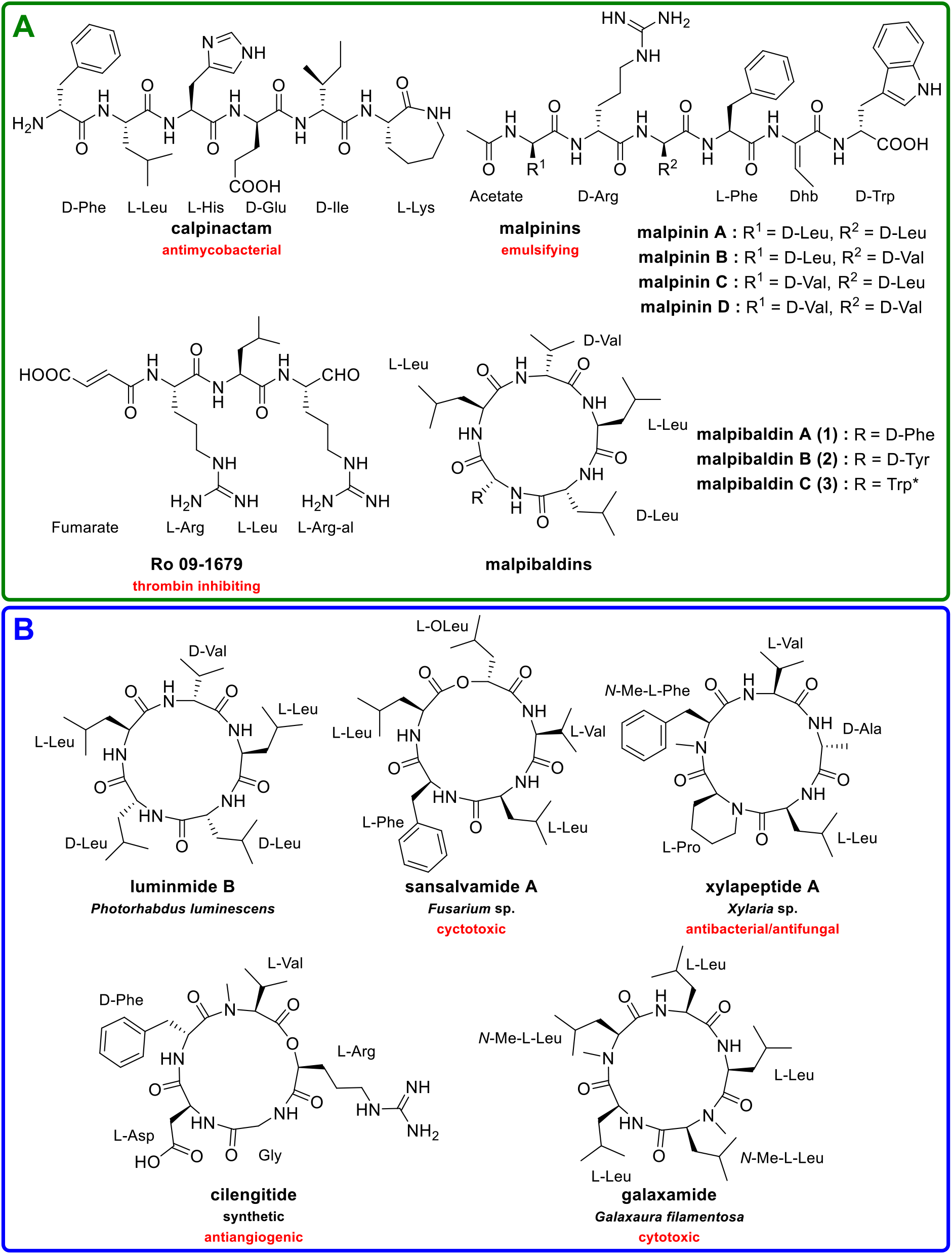
Linear and cyclic oligopeptides from *M. alpina* and other resources. (**A**) Oligopeptides isolated from *Mortierella alpina* (15–18). (**B**) Malpibaldin-related cyclopentapeptides from other sources. (19, 21-24). *Configuration of the Trp moiety was not determined (17). Dhb, dehydrobutyrine, L -Arg-al, l-arginal.

The biosynthetic origin of all above-mentioned peptides from *M. alpina* has never been illuminated. The publicly available 38.38 Mb genome of *M. alpina* (26) harbors 22 genes encoding polymodular NRPS or monomodular NRPS-like proteins indicating its high biosynthetic potential (7). However, the frequent occurrence of d-amino acids in peptides of *Mortierellales* is biosynthetically surprising, since mainly bacteria (27), but rarely fungi (28–30), use d-amino acids as building blocks for NRP synthesis. Investigations on the mucoromycete *Rhizopus microsporus* revealed that its d-amino acid-containing peptides rhizonin and heptarhizin are not produced by the fungus but by its in-host endosymbiotic proteobacterium *Paraburkholderia rhizoxinica* (31, 32). Similarly, *Burkholderia*-related (BRE) endosymbionts such as *Mycoavidus cysteinexigens* are frequently observed in *Mortierellales* (33, 34). In addition, non-culturable *Mycoplasma*-related obligate endosymbionts (MRE) have been identified by 16S-rDNA sequencing in *Mortierellomycotina* (35). Therefore, we considered a possible involvement of endosymbionts in the secondary metabolism of *M. alpina*.

Here, we report the biosynthesis of malpibaldins and a novel set of cyclopentapeptides, called malpicyclins A-F (**4-9**), in *M. alpina*. The latter compounds are composed of d/l-Leu, d/l-Val, d-Phe, d-Trp, d-Tyr and d-Arg and show moderate activity against gram-positive bacteria. The production of both peptide families does not rely on fungus-associated bacteria. Instead, two fungal genes encoding pentamodular NRPSs are expressed during metabolite production and their amino acid sequence similarity to bacterial NRPS suggests a genetic transfer across kingdoms. By combination of differential expression analysis, metabolite profiling, and *in-silico* bioinformatic substrate binding studies the NRPS genes *mpbA* and *mpcA* were specifically linked to the biosynthesis of malpibaldins and malpicyclins, respectively. Heterologous production of MpbA and MpcA modules in *Escherichia coli* and subsequent substrate turnover assays confirmed their activity.

## Results

### Metabolic profiling of *M. alpina* revealed the malpicyclin family

The high abundance of the previously reported small peptides (17) prompted us to screen alternative cultivation conditions to induce oligopeptide production in *M. alpina*. Cultivation in potato dextrose broth (PDB) resulted solely in the production of malpinins (17) in the mycelium according to UHPLC/MS measurements (Figure 2A). In contrast, when cultivated in lysogeny broth supplemented with 2% fructose (LB+ F), the previously reported malpibaldins A-C (**1-3**) (17) and an additional series of four masses [M+H]^+^ of unknown nature (**4-7**) were detected (Figure 2A). Upscaling of the culture to 4 L and subsequent isolation of the metabolites by semi-preparative HPLC enabled the identification of the four cyclopentapeptides malpicyclin A, B, C, and D (*m/z* 645.4074 [*M*+H]^+^, *m*/z 645.4065 [*M*+H]^+^, *m*/z 629.4343 [*M*+H]^+^, *m*/z 615.3976 [*M*+H]^+^, compounds **4-7**) by 1D and 2D NMR analysis (Figure 2B, Table 1, Tables S1-S4, Figures S1-S27). ^13^C NMR experiments and the calculated molecular formulae revealed by HR-MS suggested five carbonyl atoms and eight nitrogen atoms in **4-7** and indicate pentapeptides that include a guanidinium group from arginine. MS/MS experiments and ^1^H,^1^H COSY analysis suggested a cyclic ring structure which was confirmed by HMBC and HSQC 2D analysis experiments. Both COSY- and HMBC-data allowed the assignment of the amino acid side chains and revealed the planar structure: cyclo-(-Leu/Ile/Val-Tyr/Phe-Arg-Val-Leu-) (Figure S6). The absolute configuration of the side chains was elucidated by the advanced Marfey’s method (36, 37) (Table S5). d-Amino acids are solely incorporated at positions 1, 3, and 4 in **4-7**. Of note, besides d-Arg at position 3, l-Arg was confirmed as second building unit as evident by dual signals in Marfey’s analysis (Table S5) and the presence of two symmetric scalar couplings (δ_H_ 1.63 and δ_H_ 1.45 in malpicyclin C) in HSQC spectra (Fig. S21). In addition to **4-7**, two by-products (**8** and **9**) were detected by HR-MS (Table 1, Tables S1-S2), but were produced in insufficient amounts for NMR analysis. The MS/MS fragmentation of **8** (malpicyclin E, *m/z* 668.4228 [*M*+H]^+^) showed a similar pattern as for **4** (Figure S5) and suggested a tryptophan as aromatic amino acid at the 3^rd^ position (Table 1). According to HR-MS data, **9** (malpicyclin F, *m/z* 631.3959 [M+H]^+^) is probably identical to the arginine-containing cyclopeptide MBJ-0174, previously isolated from *Mortierella alpina* strain f28740 (18).

**Figure 2.**
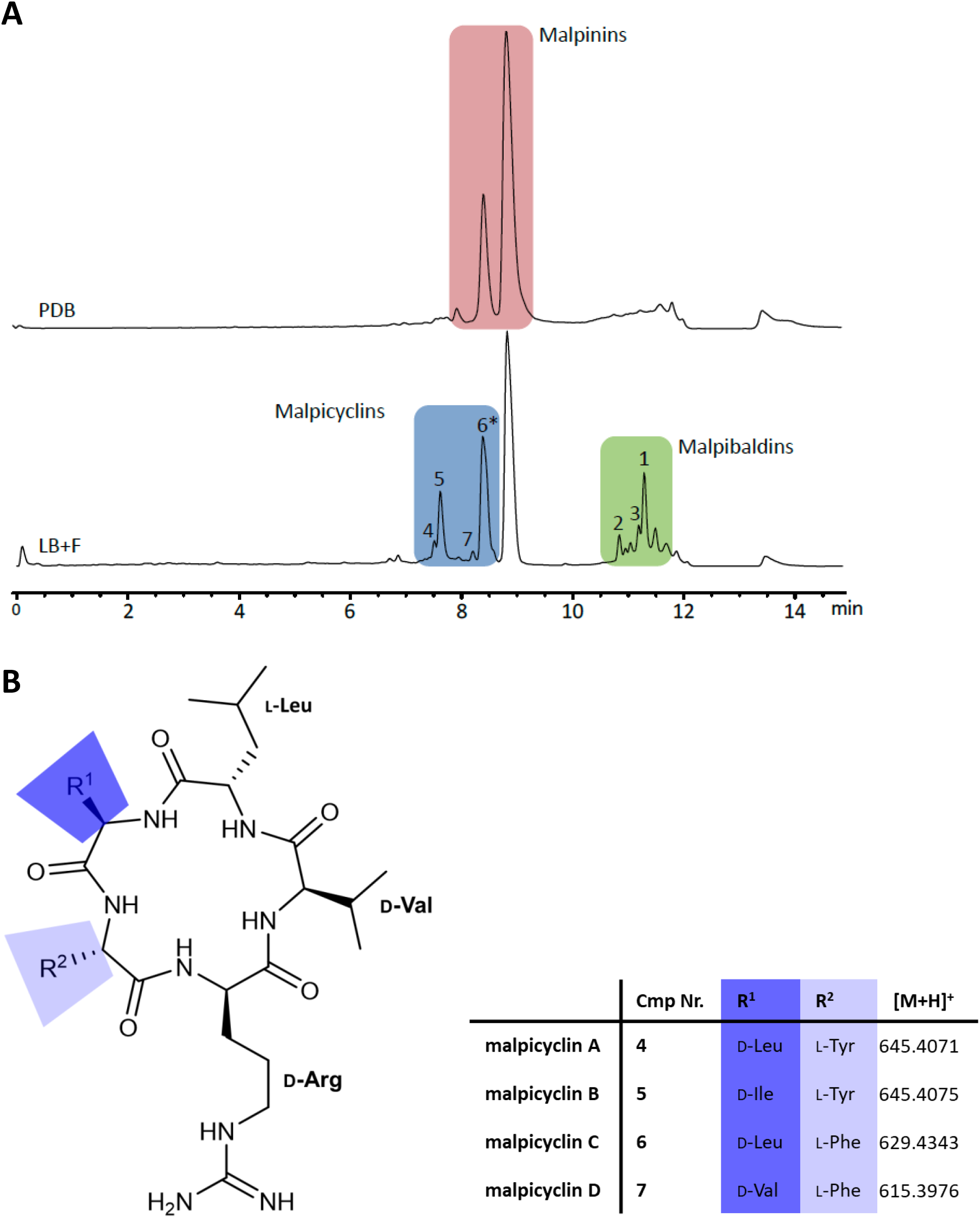
Identification of malpicyclins from *M. alpina* extracts. **A.** UHPLC-MS profile of mycelial crude extracts from *M. alpina* after cultivation in potato dextrose broth (PDB) or modified lysogeny broth (LB+F, LB medium supplemented with 2% fructose) for 7 days (TIC). **B.** Scheme of the chemical structure of malpicyclins A-D (**4-7**). * Signal overlap: Malpicyclin C (**6**) coelutes with malpinin B.

**Table 1.** HR-ESI-MS data of isolated cyclopentapeptides from *M. alpina* ATCC32222. *Absolute configuration of amino acids was not determined.

| no. | compound | m/z [M+H]<sup>+</sup> | Planar structure | Ref. |
| --- | --- | --- | --- | --- |
| 1 | malpibaldin A | 586.3963 | cyclo(-L-Leu-D-Leu-D-Phe-L-Leu-D-Val-) | (17) |
| 2 | malpibaldin B | 625.4062 | cyclo(-L-Leu-D-Leu-D-Trp-L-Leu-D-Val-) | (17) |
| 3 | malpibaldin C* | 602.3912 | cyclo(-Leu-Leu-Trp-Leu-Val-) | (17) |
| 4 | malpicyclin A (plactin B) | 645.4074 | cyclo(-D-Leu-L-Tyr-D-Arg-D-Val-L-Leu-) | (38, 39, 80) |
| 5 | malpicyclin B | 645.4065 | cyclo(-D-Ile-L-Tyr-D-Arg-D-Val-L-Leu-) | This study. |
| 6 | malpicyclin C (plactin D) | 629.4343 | cyclo(-D-Leu-L-Phe-D-Arg-D-Val-L-Leu-) | (38, 39, 80) |
| 7 | malpicyclin D | 615.3976 | cyclo(-D-Val-L-Phe-D-Arg-D-Val-L-Leu-) | This study |
| 8 | malpicyclin E* | 668.4228 | cyclo(-Leu-Trp-Arg-Val-Leu-) | This study. |
| 9 | malpicyclin F (MBJ-0174) | 631.3959 | cyclo(-D-Val-L-Tyr-D-Arg-D-Val-L-Leu-) | (18) |

Antimicrobial testings of **4-7** revealed a moderate antibacterial activity against gram-positive bacteria with MIC values ranging between 97.3 to 357.8 μM, while gram-negative representatives or fungi were not affected (Table 2, Figure S28 and S29). Of note, **4** and **6** are structurally identical to the cyclopeptides plactin B and plactin D that has formerly been isolated from an unspecified fungal strain F165 (38). Plactin D showed blood plasma dependent fibrinolytic activity by enhancing the prothrombin protease activity (39). Since amphiphilic malpinins have been described as surface-active metabolites, we also checked for tenside properties of malpicyclins by the ring tear-off method (17). Indeed, malpicyclins demonstrated a biphasic profile with a fast decrease of the surface tension up to a concentration of 62.5 μg/mL followed by a slower decrease (Figure S30). However, the calculated CMC of 93.9 μg/mL (147 μM) is 10-fold higher than that of malpinins (17) suggesting only a moderate contribution to *Mortierella*’s biosurfactant activities.

**Table 2.** Minimal inhibitory concentration (MIC_50_) of malpicyclins A-D tested against *Bacillus subtilis* and *Escherichia coli*. iprofloxacin served as positive control.

| compound | MIC<sub>50</sub> (μM) |  |
| --- | --- | --- |
|  | *B. subtilis* | *E. coli* |
| 4 | 181.1 ± 89.6 | > 1000 |
| 5 | 97.3 ± 16.5 | > 1000 |
| 6 | 357.8 ± 124.4 | > 1000 |
| 7 | 255.9 ± 15.2 | > 1000 |
| ciprofloxacin | 0.022 ± 0.002 | 0.101 ± 0.003 |

### Malpicyclins and malpibaldins are fungal products

In 1984, the macrolactone antibiotic rhizoxin was isolated from the zygomycete *Rhizopus microsporus* and was seemingly of fungal origin (40). However, 23 years later, Hertweck’s group demonstrated in pioneering work, that the fungus harbors the endosymbiotic β-proteobacterium *P. rhizoxinica* that produces this macrocyclic polyketide (41). Analogously, *Mortierella elongata* can be infected by the endobacterium *Mycoavidus cysteinexigens*, whose genome encodes at least three NRPS while the host genome does not (34). Indeed, a screening approach of 30 different *Mortierella* isolates revealed that 13% were infected by *Mycoavidus-*related endosymbionts (MRE) (34). To investigate whether cyclopentapeptides in *M. alpina* ATCC32222 are of fungal or bacterial origin, the strain was repeatedly treated with an antibiotic cocktail known to cure potential infected hyphae (41). However, metabolite production - represented by quantifying the major compound of each metabolite class (malpicyclin C (**6**) and malpibaldin A (**1**)) - was not affected by the antibiotic treatment, suggesting that the metabolite is produced by the fungus (Figure S31). Moreover, a subsequent PCR-amplification of bacterial 16S-rDNA from treated and untreated fungal mycelium of *M. alpina* failed and pointed to the absence of endobacteria (Figure S32). Both experiments indicate, that i) *M. alpina* isolate ATCC32222 does not harbor endobacteria and that ii) the isolated metabolites are rather fungal than bacterial products.

### Genome mining of *M. alpina* revealed the identification of cyclopeptide synthetases

A whole-genome survey of *M. alpina* (26) using ANTISMASH (42) revealed 22 genes encoding NRPS and NRPS-like proteins in *M. alpina*. According to the molecular structure and d-amino acid distribution in malpicyclins and malpibaldins, two five-module NRPS with at least three epimerization domains (E) or dual epimerase/condensation domains (E/C) are required. Indeed, two candidate NRPS genes (*mpcA* and *mpbA*) were identified in the genome (Figure 3A). The potential NRPS gene product MpcA (5,532 aa) comprises a pentamodular NRPS with a scaffold C_s_-A-T-E/C-A-T-C-A-T-E/C-A-T-E/C-A-T-TE which includes an expected pattern of bacterial-like dual E/C domains (in module 2, 4, and 5) as required for malpicyclin biosynthesis. The high similarity to bacterial domains facilitated the prediction of the *M. alpina* A domain specificities which can be a difficult task for fungal domains. Analysis of putative substrate preference of the A domains by alignment with the GrsA A domain from *Aneurinibacillus migulanus* (43) suggested acceptance of hydrophobic amino acid substrates in all A domains of MpcA with exception of domain A3 (Table 3, Table S6). Here, hydrophilic residues (Ser, Thr) are found at position 278 and 330 (GrsA numbering) of the NRPS code and, moreover, an Asp at position 331 might ion-pair with cationic amino acids such as Arg (44, 45) which is abundant in all malpicyclins (**4-7**). Hence, we hypothesized that a positively charged amino acid would be accepted by the MpcA A3 domain.

**Table 3.** NRPS codes for adenylating domains of the *Mortierella* NRPS (MpcA and MpbA) and related biochemically investigated bacterial and fungal NRPS modules. The proposed activated substrates are listed in column 3. The residues are mapped relative to *Aneurinibacillus migulanus* (former *Brevibacillus brevis)* GrsA-A numbering (81). NRPS codes were partially extracted from Bian *et al.* (82). Amino acid residues in the NRPS code are highlighted according to their physicochemical properties: acidic (red), small/hydrophobic (grey), aromatic/hydrophobic (amber), hydrophilic (green), and basic (blue). *Organisms are highlighted according to their phylogenetic origin: basal fungi (blue), higher fungi (brown), and bacteria (green). A detailed alignment is provided as supplementary table (Table S6).

| Protein domain | Organism* | substrate | Residue position according to GrsA Phe numbering |  |  |  |  |  |  |  |  |  |
| --- | --- | --- | --- | --- | --- | --- | --- | --- | --- | --- | --- | --- |
|  |  |  | 235 | 236 | 239 | 278 | 299 | 301 | 322 | 330 | 331 | 527 |
| **MpcA_A2** | *M. alpina* | Tyr/Phe | D | P | F | T | M | G | A | V | V | K |
| **MpbA_A3** | *M. alpina* | Trp/Tyr/Phe | D | P | F | V | M | G | G | T | V | K |
| Plu3263_A3 | *P. luminescens* | Phe | D | A | W | C | I | A | A | V | C | K |
| BacC_A2 | *B. subtilis* | Phe | D | A | F | T | V | A | A | V | C | K |
| CepA_A1 | *A. orientalis* | Tyr | D | A | S | T | V | A | A | V | C | K |
| GliP_A1 | *A. fumigatus* | Phe | D | G | S | I | L | G | A | C | A | K |
| **MpbA_A1/** | *M. alpina* | Val | D | A | F | W | L | G | G | T | F | K |
| **MpcA_A4** |  |  |  |  |  |  |  |  |  |  |  |  |
| **MpcA_A1** | *M. alpina* | Leu/Ile/Val | D | A | F | F | I | G | A | M | L | K |
| **MpbA_A2** | *M. alpina* | Leu | D | A | F | F | I | G | A | M | V | K |
| **MpbA_A4/A5** | *M. alpina* | Leu | D | A | I | F | L | G | A | T | I | K |
| **MpcA_A5** | *M. alpina* | Leu | D | A | M | F | I | G | G | T | I | K |
| Plu3263_A2/A4/A5 | *P. luminescens* | Leu | D | A | W | C | I | G | A | V | C | K |
| GrsB_A1 | *A. migulanus* | Val | D | A | F | W | I | G | G | T | F | K |
| CssA_A9 | *T. inflatum* | Val | D | A | W | M | F | A | A | V | L | K |
| **MpcA_A3** | *M. alpina* | Arg | D | A | A | S | V | G | A | T | D | K |
| SyrE_A5 | *P. syringae* | Arg | D | V | A | D | V | C | A | I | D | K |
| McyC_A1 | *M. aeruginosa* | Arg | D | V | W | T | I | G | A | V | D | K |
| PpzA-1_A2 | *E. festucae* | Arg | D | V | S | D | T | G | A | P | T | K |

**Figure 3.**
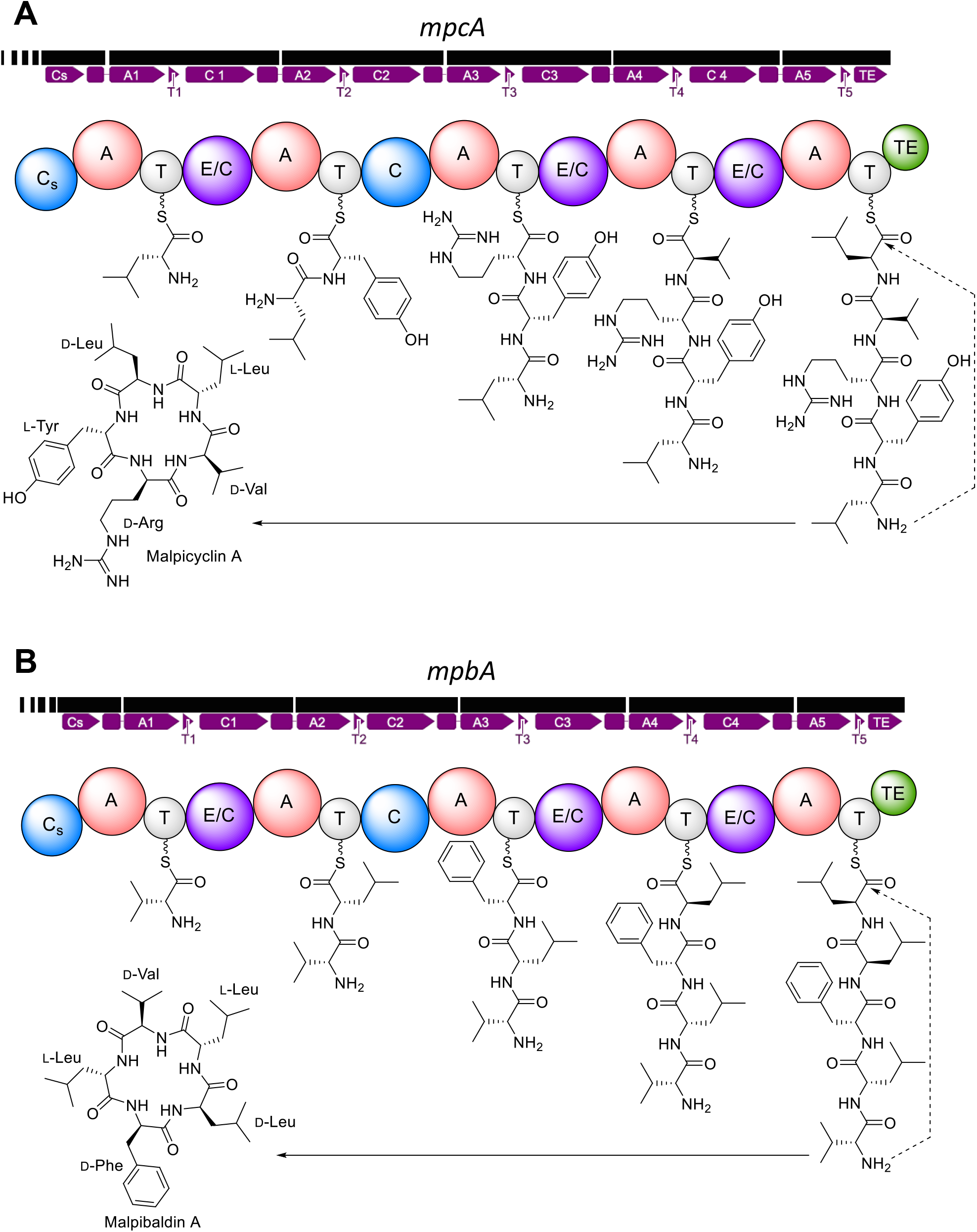
Proposed biosynthesis of malpicyclin A and malpibaldin A. **A.** Gene structure of *mpcA* and domain structure of its encoded malpicyclin synthetase MpcA. **B.** Gene structure of *mpbA* and domain structure of its encoded malpibaldin synthetase MpbA. A, adenylating domain; C condensation domain of the canonical ^L^C_L_ type; C/E, dual condensation/epimerization domain; C_s_, truncated starter condensation domain; T, thiolation domain; TE, thioesterase domain. Either gene (bold black lines) of is interrupted by nine introns. Vertical white lines represent intron, black squares exons.

The second candidate gene encodes a NRPS (MpbA) with an identical domain pattern and a size (5,541 aa) similar to MpcA (73.6% aa sequence identity). However, *in silico* substrate specificity analysis revealed exclusively hydrophobic binding pockets in all five A domains as expected for the malpibaldin building blocks (Table 3). Both MpbA A3 and MpbA A2 share a similar NRPS code suggesting that both domains accept the same - probably aromatic - substrates. MpbA is structurally related to the bacterial luminmide B synthetase Plu3263 from *P. luminescens* (50.2% identity; 65.9% similarity) and both NRPS may produce highly similar cyclopentapeptides NRP (Figures 1 and 2). However, the specificity codes of the A domains are not identical (Table 3) suggesting that zygomycetous NRPS use a dissimilar code.

In contrast to Plu3263, both MpcA and MpbA contain a starter condensation (C_S_) domain-like *N*-terminus (252 or 247 aa, respectively) known to transfer β-hydroxy-carboxylic acid residues from acyl-CoA donors to the *N*-terminus of bacterial lipopeptides (46). However, the domains are *N*-terminally truncated and the tandem histidine motifs (HH) responsible for the deprotonation of the substrate prior to the condensation step (47) is missing in the active sites of both domains. Taken together, the proposed domain structure and distribution of dual E/C domains fulfilled the requirement for incorporation of d-amino acids at the expected positions in malpicyclins and malpibaldins, and prompted us to study the malpicyclin synthetase gene (*mpcA*) and malpibaldin synthetase gene (*mpbA*) in detail.

### Fungal NRPS genes *mpcA* and *mpbA* are co-expressed in presence of fructose

To confirm active transcription of the NRPS genes, qRT-PCR expression experiments were performed. The fungus was cultivated in media to induce (LB+F) or repress (PDB) cyclopeptide production (Figures 2A and 4A). Gene expression was determined after 48 hr and 4 days of cultivation and data were normalized against expression profiles derived from freshly germinated *M. alpina* mycelium, that do not produce secondary metabolites (data not shown). Neither NRPS gene *mpcA* nor *mpbA* was expressed in PDB medium, but expression was induced 5.4- or 8.2-fold, respectively, after 48 hr cultivation in LB medium compared to PDB medium. These results matched the observed enrichment of malpibaldin A (**7**) and malpicyclin C (**3**) in mycelial metabolite extracts obtained from cultivation in LB medium (Figure 4B). In contrast, negligible amounts of both metabolites were found in mycelium after cultivation in non-inducing PDB medium.

**Figure 4.**
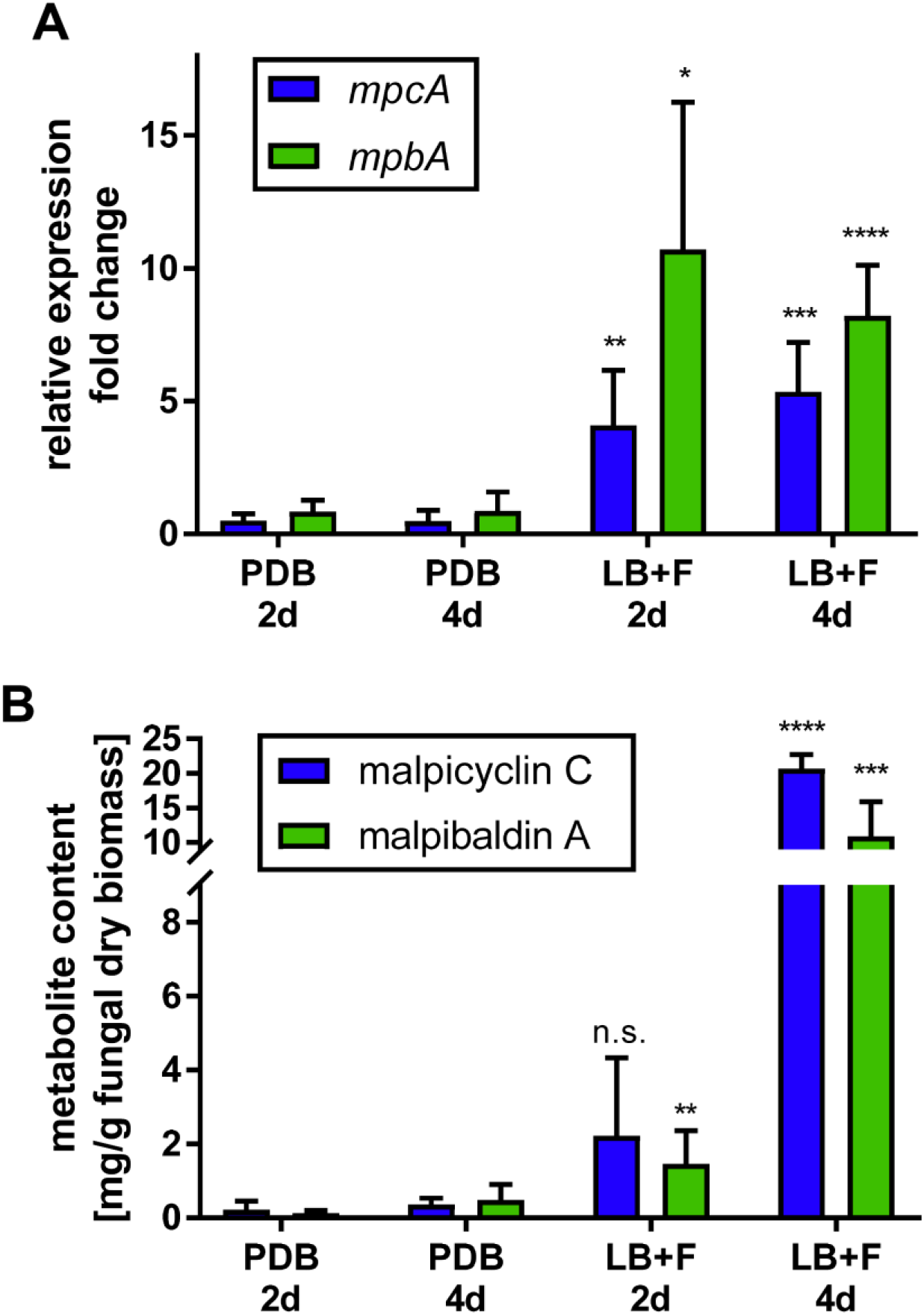
Gene expression of malpicyclin and malpibaldin synthetase genes, *mpcA* and *mpbA*, and metabolite production. *M. alpina* was cultivated under non-inducing (potato dextrose broth, PDB) and inducing conditions (lysogeny broth supplemented with 2% fructose, LB+F) up to 4 days. Expression analysis and metabolite quantification by UHPLC-MS was carried out 2 days and 4 days p. i. **A.** Expression analysis of *mpcA* and *mpbA*. Gene expression was normalized against the housekeeping genes *actA*, encoding α-actin, and *gpdA*, encoding the glycerinaldehyde-3-phosphate dehydrogenase. cDNA from freshly germinated mitospores (in PDB) served as reference (set to 1). **B.** Metabolite quantification of the malpicyclin C (**6**) and malpibaldin A (**1**). Representative of each series, the amount of the most abundant metabolites (malpicyclin C (**6**) and malpibaldin A (**1**)) was determined by UHPLC-MS mass chromatograms (EIC) from *M. alpina* mycelium. Statistical significance compared to non-induced control (PDB, 2d): n.s, not significant; * p ≤ 0.05; ** p ≤ 0.01; *** p ≤ 0.001 (paired student’s t-test). Metabolite production and gene expression correlated with a Pearson correlation coefficient r = 0.84 (p = 0.004) for malpicyclin C/*mpcA* and r = 0.72 (p=0.02) for malpibaldin A/*mpbA* at day 4.

### Adenylation domains of MpcA and MpbA activate l-amino acids

Unlike *Aspergilli*, *Mortierellaceae* are hardly genetically tractable and targeted gene deletion occurs with very low frequency and genomic stability (48, 49). Hence, we verified the substrate specificity of the most characteristic modules (C-A-T) in both NRPS, i.e. module 3 of MpcA (MpcA-m3, 122.45 kDa) and module 3 of MpbA (MpbA-m3, 122.40 kDa), by heterologous production in *E. coli* (Figure S33). The purified double-His_6_-tagged fusion proteins (50) were subjected to specificity testing by the recently established MS/MS-based HAMA-assay (51), which detects the formation of stable amino acid hydroxamates after enzymatic adenylation by A domains. All proteinogenic l-amino acids and two of its enantiomer counterparts (d-Val and d-Phe) were tested in parallel.

For MpcA-m3, solely the corresponding l-Arg product formation was observed, confirming the findings of the NMR analysis and *in-silico* substrate prediction (Figure 5A). However, due to a lack of an authentic hydroxamate standard, incorporation of d-Arg could not be excluded. Hence, in a complementary experiment, substrate specificity was confirmed by an ATP-[^32^PP_i_]-exchange assay (52) (Figure S34, Table S7). First, pools of physiochemically similar l-amino acids were tested, followed by a subsequent determination of single amino acids in the active pools. Again, MpcA-m3 showed highest turnover with l-Arg (2.2 × 10^5^ dpm). However, 11% of ATP/^32^PP_i_ conversion was also observed for d-Arg (2 × 10^4^ dpm), suggesting that the A domain is not entirely enantioselective. Poor discrimination against d-amino acids is not surprising when the d-enantiomers are not present as competing substrates. For instance, strong side activity for d-Phe and d-homoserine (d-Hse) has been reported for usually l-amino acid activating A domains in tyrocidine and ralsolamycin biosynthesis, respectively (52, 53).

**Figure 5.**
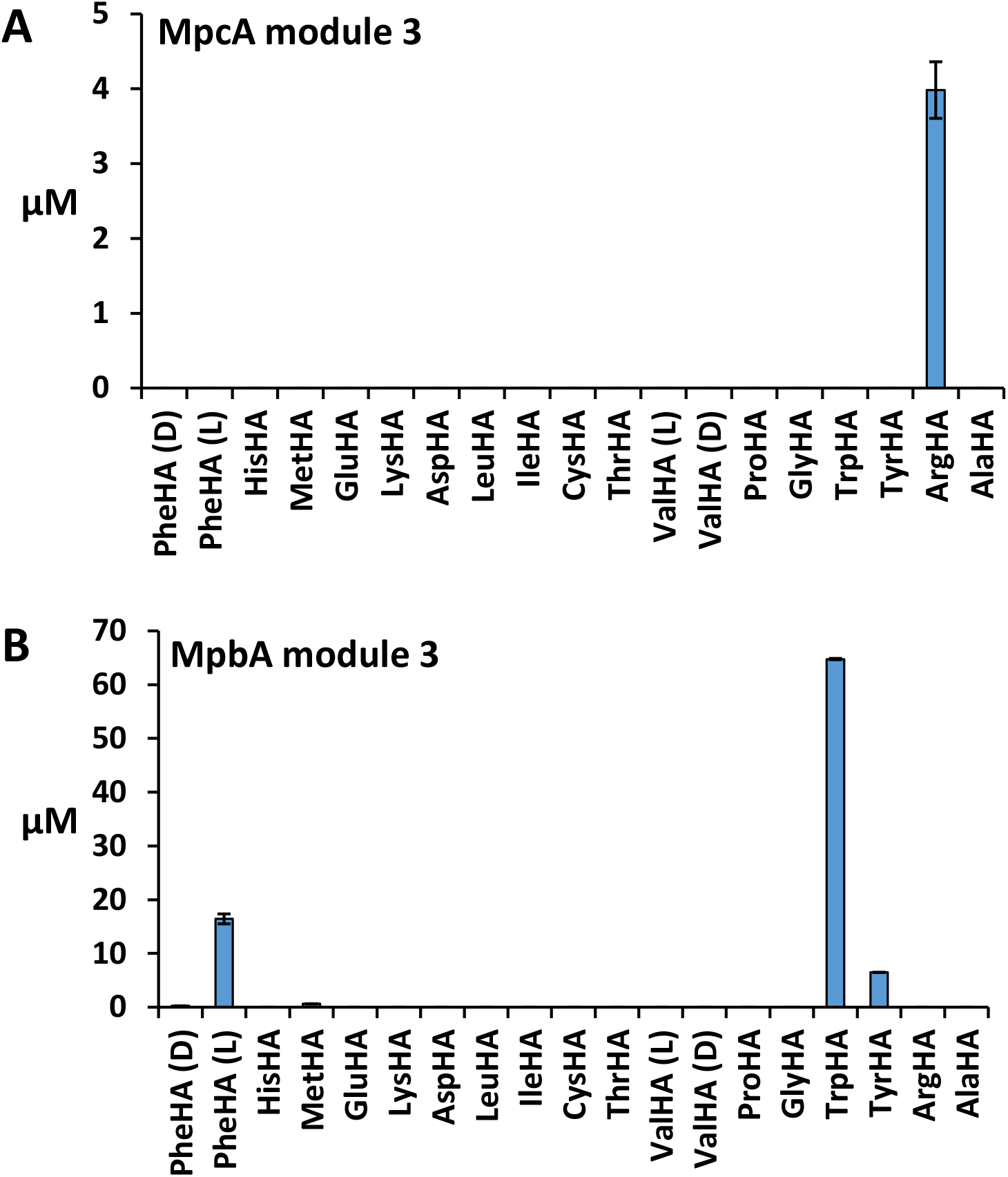
Substrate specificity testing of NRPS modules by the multiplexed hydroxamate assay (HAMA). Modules 3 (C-A-T) of MpcA (**A**) and of MpbA (**B**) were separately tested. Substrates were all proteinogenic amino acids (except l-Asn, l-Gln, l-Ser), d-Phe and d-Val. Amino acyl hydroxamates were quantified using HAMA (n = 3).

The HAMA-assay of MpbA-m3 revealed a promiscuous acceptance of aromatic amino acids, with preference l-Trp>l-Phe>l-Tyr (Figure 5B). These side chain identities perfectly match residues found in malpibaldin C, A and B, respectively, but are not in line with the preference for product formation. In *M. alpina*, malpibaldin A (l-Phe derivative) is the major compound (17). Such discrepancies between adenylation preference and relative product formation rate may be caused by side-chain specificity of downstream reaction steps or differences in intracellular amino acid availability (51). Promiscuous adenylation leading to the parallel production of multiple products from one NRPS assembly line has been well studied in cyanobacterial enzymes and may be an important springboard for evolutionary diversification (45, 54).

From this knowledge, we predicted the biosynthesis of malpicyclins and malpibaldins by successive activation and, if required, racemization of l-amino acids, which are subsequently condensed to give a nascent, linear pentapeptide tethered to the T-domains of the enzymes (Figure 3). The final cyclization is probably catalyzed in *cis* by the *C*-terminal type I thioesterase (TE) domain, which has been demonstrated for several cyclopeptides from bacteria (52, 55, 56).

### NRPS genes of *M. alpina* are of (endo)bacterial origin

Since no zygomycetous NRPS has been identified before, we were interested in about the evolutionary origin of the genes and enzyms. A phylogenetic analysis of the extracted A domains from both *M. alpina* NRPS proteins and from verified fungal and bacterial NRPS, NRPS-like and PKS-NRPS hybrid proteins was performed. Surprisingly, the zygomycetous A domains clustered in a monophylum with bacterial - but not with fungal - A domains independent of substrate specificity (Figure 6, Table S8). Moreover, the A domains share high similarity to A domains from NRPS of endobacteria such as *Paraburkholderia* sp. or *Mycoavidus* sp., known to infect zygomycetes, but are more distantly related to other gram-negative (*Pseudomonas* or *Ralstonia*) or gram-positive representatives *(Streptomycetes* sp.). Hence, both *Mortierella* NRPS are most likely of (endo)bacterial but not fungal origin and may have been evolved independently of the fungal counterparts.

**Figure 6.**
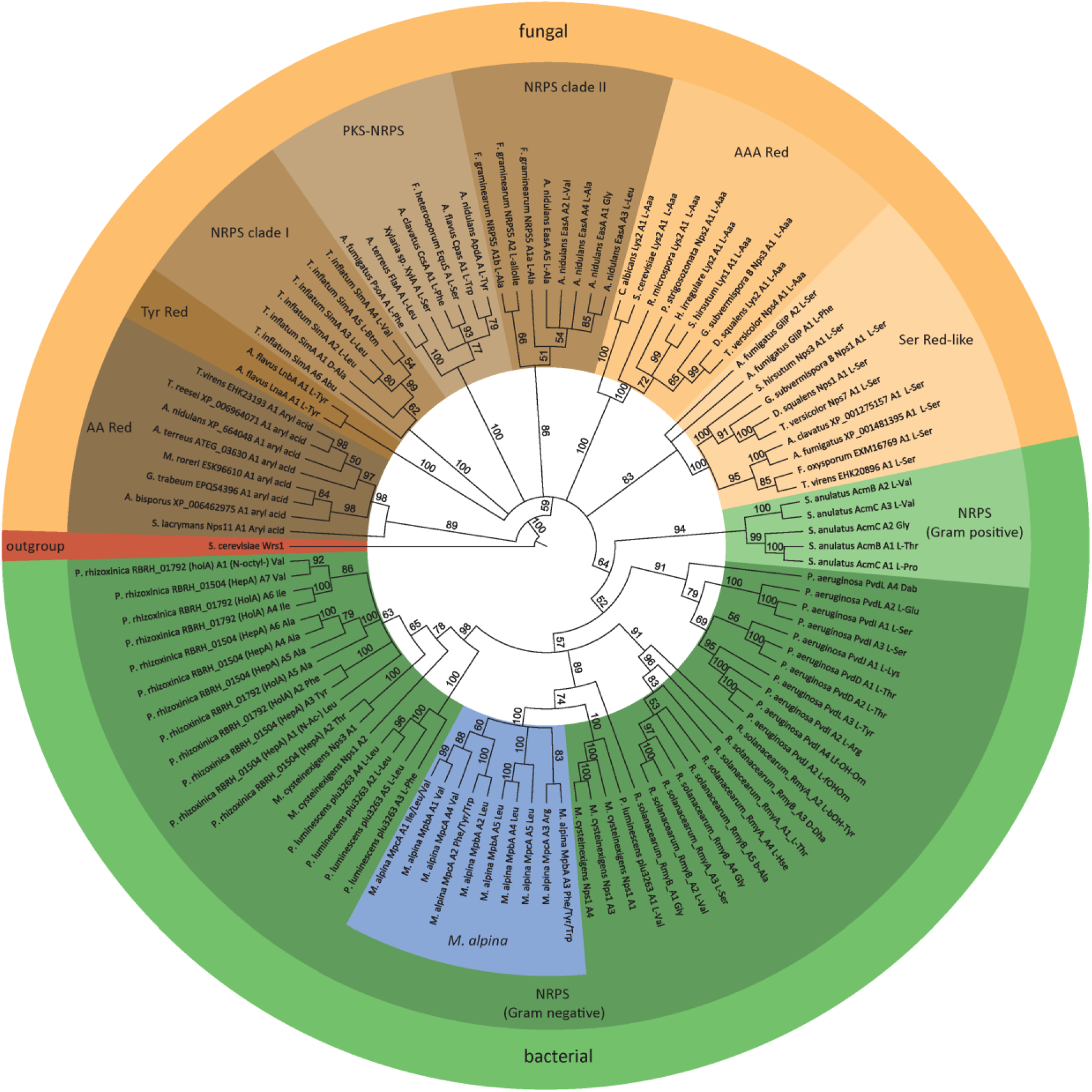
Phylogenetic analysis of A domains from *M. alpina* and other fungal or bacterial representatives. The A domains were extracted from NRPS, NRPS-like proteins and PKS/NRPS hybrids from *M. alpina* (blue), (endo-)bacteria (green) and higher fungi (taupe/amber) (refer to Table S8). The A domain of the cytoplasmatic tryptophanyl-tRNA synthetase Wrs1 from *Saccharomyces cerevisiae* served as outgroup (red). Note that the “zygomycetous A domains” from *M. alpina* cluster together with bacterial A domains. The percentual bootstrap support is labeled next to the branches. AAA red, α-2-aminoadipate reductase; AA Red, aryl acid reductase; NRPS, nonribosomal peptide synthetase; PKS-NRPS, polyketide synthase-nonribosomal peptide synthetase hybrid; Ser Red, serine reductase; Tyr Red, tyrosine reductase.

A phylogenetic analysis based on the more conserved condensation (C) domains showed a similar outcome (Figures S35 and S36, Tables S9 and S10). Most interestingly, C domains from *M. alpina* NRPS fall into three groups: ^L^C_L_, common C domains condensing l-amino acids, C_s_, *N*-terminal starter condensation domains, and dual E/C domains, epimerizing enzyme-tethered l-amino acids to its d-enantiomer prior to condensation. The canonical ^L^C_L_ domains from *Mortierella* cluster both with bacterial and fungal ^L^C_L_ domains (Figures S35 and S36). In addition, the truncated C_s_ domains have been most likely evolved from *Mortierella* ^L^C_L_ domains and show no similarity to acyl-transferring C_s_ domains found in bacterial lipopeptide NRPSs (Figures S35). In contrast, the six dual C/E domains from MpcA and MpbA cluster with those of endobacteria such as *Paraburkholderia* spec. (Figure S35), but have no close counterpart among fungal C domains (Figure S36). To date, dual E/C domains have been solely found in bacteria but not in fungi. Indeed, the zygomycetous E/C domains occupy in a unique, separate C domain clade among the fungal kingdom.

Taking all these findings together, we postulate a probable horizontal gene transfer (HGT) from an unknown bacterial endosymbiont to the *Mortierella* host. However, the GC content of *mpcA* (55.0%) and *mpbA* (54.6%) is similar to that of the *M. alpina* genome (51.8%) and the codon usage of their coding sequences is nearly identical to that of the housekeeping genes from *M. alpina* (Figure S37). Both findings suggested an early HGT event in *Mortierella*.

## Discussion

Zygomycetes of the order *Mortierellales* are an established resource for enzymes in industrial detergent manufacturing or for polyunsaturated fatty acids in food industry. Recent publications revealed highly bioactive compounds such as the antiplasmodial cycloheptapeptide mortiamides A (57) or the surface active hexapeptide malpinin A (17). Hence, this fungal order seems to be a prolific resource of pharmaceutically relevant natural products, too. However, the biosynthetic basis of the peptide compounds from zygomycetes has never been investigated. For the first time, we gave evidence that secondary metabolite genes are actively transcribed in zygomycetes and encode functional enzymes, called “zygomycetous NRPS”.

The zygomycetous NRPSs MpcA and MpbA combine properties of both bacterial and fungal NRPS. A bacterial origin of the NRPS genes is likely since they encode A and C domains, that exclusively cluster with bacterial representatives. Most of the as yet known NRP from *Mortierella* species use d-amino acids as building blocks. To incorporate d-amino acids into the NRP backbone, fungi require a separate, pre-NRPS acting amino acid racemase as shown for the biosynthesis routes of cyclosporine in *Tolypocladium niveum* or HC-toxin in *Cochliobolus carbonum* (28, 58). Alternatively, fungal NRPS possess a dedicated epimerase domain *N*-terminally located to a ^D^C_L_ domain that solely accept d-amino acids as substrate, i.e. in the fusaoctaxin A synthetase NRPS 5 from *Fusarium graminearum*. (29) In contrast, zygomycetous NRPS use E/C domains that have not been described in fungi before. Bacterial E/C domains are characterized by two consecutive histidine repeats (HH) in condensation domain motifs C1 (HH[I/L]xxxxGD) and C3 (HHxxxGDH) which are required for epimerization and condensation, respectively (59). Similar catalytic HH motifs are present in the *Mortierella* dual E/C domains (C1: HH[M/L][M/L]A[T/A]EGD and C3: HH[I/L][IV][G/I]DH) suggesting a functional racemization.

Another distinctive feature of the zygomycetous NRPS is the *N*-terminal truncated starter C-domain (C_s_) in both MpcA and MpbA. *Mortierella* C_s_ domains do not cluster with β-hydroxy acyl-transferring C_s_ domains from lipopeptide producing NRPS from bacteria (46). Instead, our phylogenetic analysis indicated that these domains have been evolved from the *Mortierella* canonical ^L^C_L_ domains. Truncated C_s_ domains with unknown function were also found in fungal NRPS such as the cyclosporin synthetase SimA from *T. inflatum* (60), the tryptoquialanine synthetase TquA from *Penicillium aethiopicum* (61), and the pyrrolopyrazine synthetase PpzA-1 from *Epicholoe festucae* (62) and are thought to be an evolutionary relic required to maintain A domain stability (63). In bacterial macrocyclic-producing NRPS, the final peptide is cyclized by a terminal *cis*-acting type I TE domain, which is often replaced by a specialized cyclase-like C domain (C_T_) in fungi (30). Both domains preferably catalyze a head-to-tail macrolactamization of peptides with d- and l-configured residues at the N- and C-terminus, respectively (64). Hence, the presence of C-terminal TE-domains in MpcA and MpbA and the presence of a dual E/C domain at their N-termini points at a bacterial-like cyclisation mechanism in zygomycetous NRPS.

Surprisingly, based on the amino acid sequence, MpbA is highly related to bacterial NRPS’s, such as the luminmide B synthetase Plu3263 from *P. luminescens.* Both NRPS’s produce highly similar NRP with identical amino acid configuration (24). The A domains of Plu3263 are extraordinary flexible and accept a variety of (non)-proteinogenic amino acid substrates resulting in the production of novel luminmide variants (luminmide C to I) by simple substrate feeding (65). Similarly, in *M. alpina* quantity and quality of malpibaldins and malpinins is dependent on the amino acids supplied in the medium (17), which was confirmed by the promiscuity of the MpbA A3 domain accepting at least three aromatic substrates. Interestingly, the NRPS code of the zygomycetous A domains show higher similarity to bacterial than to fungal A domains. E.g. the arginine-activating A3 domain of the MpcA comprise a guanidinium-stabilizing aspartate residue which is also present in the arginine-adenylating A domains of the bacterial NRPS from *Pseudomonas syringae* pv. *syringae* (66) or *Microcystis aeruginosa* (54) but is absent in fungal NRPS such as from the ascomycete *E. festucae* (62).

Taking all these finding together and considering the fact that *Mortierella* strains are frequently infected with NRPS-encoding proteobacteria from the genus *Mycoavidus* ssp. (35), a horizontal gene transfer (HGT) from an (endo)bacterial symbiont into the fungal host is highly plausible. A similar phenomenon is postulated for the genes for β-lactam antibiotics: The responsible ACVS-like NRPS genes from *Aspergillus nidulans* and *Penicillium chrysogenum* have most likely arisen from *Lysobacter*-like or *Streptomyces*-like bacterial ancestors (67–69). In phylogenetic analysis, A domains of fungal ACVS-NRPS cluster with bacterial representatives into one unique monophyletic group (70), similarly as MpcA and MpbA with endobacterial NRPS in this study. However, neither codon usage nor GC content of the *Mortierella* NRPS genes match that of the endosymbiotic *Mycoavidus* genes but are highly similar to that of *Mortierella* housekeeping genes. As expected for fungal genes, both *mpcA* and *mpbA* contain introns of an average size of 102 bp, indicating an comparably early gene transfer during evolution of *M. alpina* and a subsequent adaptation of the genes to the requirements of the eukaryotic mRNA processing (71). Recently, HGT events have been postulated as the main driver of secondary metabolism diversity in the zygomycetous genus *Basidiobolus* as supported by the phylogenetic reconstructions of NRPS gene clusters with bacterial homologs (72). *Basidiobolus* spec. are common inhabitants of the amphibian gut and, similar to *Mortierella* spec., live in close association with proteobacteria (73).

The ecological function of zygomycetous NRPs is still to be deciphered. However, from a pharmaceutical angle, malpicyclins A and C are structurally identical to the cyclopentapeptide plactin B and D, respectively, which were previously isolated from an uncharacterized fungus (38, 74) and exhibit fibrinolytic activities by elevating the activity of cellular urokinase-type plasminogen activator (39). The anticoagulating effects of plactins and derived cyclopentapeptides such as malformin A_1_ are currently under investigation in treatment of thrombic disorders (75). Hence, *M. alpina* is not only of nutritional benefit by production of polyunsaturated fatty acids, but also of pharmaceutical interest as producer of bioactive natural products. Moreover, the abundance of natural products in *M. alpina* under certain growth conditions raises safety concerns regarding the biotechnological use of this strain.

In sum, this report disproves a long-standing dogma of a marginal secondary metabolism in zygomycetes and, instead, establishes *Mortierellales* as a promising, previously overlooked reservoir for bioactive metabolites.

## Material and Methods

### Organisms and culture maintenance

The fungal strain *Mortierella alpina* ATCC32222 was purchased from the American Type Culture Collection (ATCC). Cultures were maintained on MEP agar plates (30 g/L malt extract, 3 g/L soy peptone, 18 g/L agar) for 7 days at 25 °C. For antibiotic treatment of *M. alpina* and subsequent analyses (16S-rDNA detection, metabolite quantification), refer to the Supplemental Experimental Procedures. *Bacillus subtilis subsp. subtilis* 168 (DSM 23778) and *Escherichia coli* DSM 498 were maintained on LB agar plates at 37°C. *E. coli* XL1-blue (Agilent) and *E. coli* KRX (Promega) were used for plasmid propagation and for protein production, respectively, and were maintained in LB (5 g/L yeast extract, 10 g/L tryptone, 10 g/L sodium chloride) medium supplemented with 50 μg/ml carbenicillin or 100 μg/ml kanamycin (both Sigma Aldrich), if applicable.

### Chemical analysis and metabolite structure elucidation

#### General

All chromatographic methods are summarized in Table S13. UHPLC-MS measurements of **1-9** were carried out on an Agilent 1290 infinity II UHPLC coupled with an Agilent 6130 single quadrupole mass spectrometer (positive ionization mode) using method A and B (Table S13) for routine metabolite detection and for metabolite quantification. High-resolution mass spectra and MS/MS fragmentation pattern of **1-8** were recorded using a Q Exactive Plus mass spectrometer (Thermo Scientific). Chromatography for determination of amino acyl hydroxamate was performed on a Waters ACQUITY H-class UPLC system coupled to a Xevo TQ-S micro (Waters) tandem quadrupole instrument with ESI ionisation source in positive ion mode (desolvation gas: N_2_, collision gas: Ar, capillary voltage 1.5 kV, cone voltage 65 V, desolvation temperature 500°C, desolvation gas flow 1000 L/h). NMR spectra were recorded on a Bruker Avance III 600 MHz spectrometer at 300 K using DMSO as solvent and internal standard. Peaks were adjusted to δ_H_ 2.49 ppm and δ_C_ 39.5 ppm.

#### Metabolite isolation, structure elucidation and antibiotic testings

Eight flasks with 500 mL LB medium amended with 2 % (w/v) fructose were inoculated with six agar blocks (2 × 2 mm) of *M. alpina* grown on MEP agar. After seven days of cultivation at 160 rpm at 25°C, mycelium was harvested, resuspended in 1 L methanol/butanol/DMSO (12:12:1) and homogenized using a blender (Ultra turrax, IKA). The extract was filtered, and extraction of the fungal biomass was repeated. Both extracts were pooled and evaporated to dryness. The residue was resuspended in 25 mL methanol/DMSO (10:1) subjected to an Agilent Infinity 1260 preparative HPLC equipped with a Luna C_18_ column (250 × 21.2 mm, 10 μm, Phenomenex). The metabolites were separated according to method C (Table S13) (t_R_ =8-10 min). Subsequently, the compounds were purified using method D (Table S13) on an Agilent 1200 HPLC system. Purified compounds were dissolved in DMSO-*d6* for NMR analyses. Absolute configurations of amino acids in the peptides **4-7** were determined using Marfey’s method (see Supplemental Experimental Procedures and method E, Table S13). Antimicrobial activities were determined by agar diffusion plates according to a published procedure (17) and MIC analysis was carried out as described in the Supplemental Experimental Procedures.

### NRPS identification and expression analysis

The genome of *M. alpina* ATCC32222 was accessed from JGI under the GeneBank assembly accession number GCA_000240685.2. Putative NRPS genes were annotated using the fungal antiSMASH 5.0 software (42) and, if required, putative intron-exon junctions were curated manually by alignment to fungal/bacterial NRPS (Tables S8-S10). Expression primers for *mpcA* and *mpbA* as well as housekeeping genes encoding β-actin (*actB)* and the glycerinaldehyde-3-phosphate dehydrogenase (*gpdA)* were designed (cut-off PCR efficiency of at least 0.95) (Table S11). *M. alpina* was grown in LB medium amended with 2% of fructose (LB+F medium) or in potato dextrose broth (PDB, Sigma Aldrich) at 160 rpm at 25 °C for up to 4 days. RNA was extracted using the SV Total RNA Isolation System (Promega) and residual gDNA was digested with Baseline-Zero DNase (Biozym). cDNA was synthesized with the RevertAid Reverse Transcriptase (Thermo Fisher) using oligo(dT)_18_ primers. For quantitative real-time PCR (qRT-PCR), the qPCR Mix EvaGreen (BIOSELL) was used in a qPCR Cycler qTower^3^ (AnalytikJena) following the manufacturer’s instructions and qPCR protocol: initiation 95 °C, 15 min; followed by 40 amplification cycles (95 °C, 15 s; 60 °C, 20 s; 72 °C, 20 s), and final recording of a melting curve (60-95 °C). Expression data were calculated according to the ΔΔC_T_ method by Pfaffl (76) using the housekeeping genes as internal, non-regulated reference controls.

### Heterologous protein production and determination of enzymatic activity of A domains

For detailed cloning procedures and protein production protocols refer to the Supplementary Experimental Procedures and Tables S11-S12. In brief, intron-free coding sequences of MpcA module 3 (*mpcA*-M3) and MpbA module 3 (*mpbA*-M3) were amplified from cDNA and ligated into the blunt pJET1.2 vector system (Thermo) prior to final subcloning in pET28a(+) expression vectors. NRPS modules were produced in *E. coli* KRX (Promega) at 16°C using 0.1% l-rhamnose as inducer. Proteins were purified from cell-free lysate by metal ion affinity chromatography with Protino Ni-NTA Agarose (Macherey-Nagel) as matrix followed by ultrafiltration (Amicon Ultra-15 centrifugal filter units, Merck).

#### ATP-[^32^P]PP_i_ exchange assay

The assay was carried out as previously described (52, 63) using 5 nM MpcA-m3 in the 100 μL reaction mix. Initially, pools of l-amino acids were used as substrates (Table S7) followed by testing of single substrates.

#### Multiplexed hydroxamate based adenylation domain assay (HAMA)

The hydroxamate formation assay was conducted as previously described (51). In brief, the assay was carried out at room temperature in 100 μL volume containing 50 mM Tris (pH 7.6), 5 mM MgCl_2_, 150 mM hydroxylamine (pH 7.5-8, adjusted with NaOH), 5 mM ATP (A2383, Sigma), 1 mM tris(2-carboxyethyl)phosphine (TCEP) and 1 μM of enzyme. Reactions were started by adding a mix of 5 mM amino acids in 100 mM Tris (pH 8) to a final concentration of 1 mM or only buffer as a control. l-Phe, l-Val and l-Leu were distinguished from d-Phe, d-Val and l-Ile respectively by using enantiopure, deuterium labeled standards. Reactions were quenched by 10-fold dilution in acetonitrile containing 0.1 % formic acid and submitted to UPLC-MS analysis (method F, Table S13). Compounds were detected via specific mass transitions recorded in multiple reaction monitoring (MRM) mode. Data acquisition and quantitation were conducted using the MassLynx and TargetLynx software (version 4.1). Quantitation was done by external calibration with standard solutions of hydroxamates ranging from 0.0032 to 10 μM.

### Phylogenetic analysis

The genomes of the zygomycetes *M. alpina* ATCC32222, and the endofungal bacteria *Mycoavidus cysteinexigens* AG77 and *Paraburkholderia (syn. Mycetohabitans) rhizoxinica* HKI 0454^T^ were obtained from the JGI fungal genomics resource database or NCBI genome database and were subjected to a screening analysis for fungal and bacterial secondary metabolite gene clusters by the antiSMASH 5.0 software (42). The A and C domains were extracted from putative enzymes and, additionally, from experimentally proven endobacterial, bacterial, and fungal NRPS and NRPS-like enzymes (Table S8-S10). For A domain phylogeny, a total set of 108 amino acid sequences (Table S8) were aligned using the ClustaIW algorithm implemented in the Geneious 10.2.4 software. For phylogenetic analysis of C domains, altogether 225 bacterial and 90 fungal sequences of C domains of verified NRPS and NRPS-like proteins (Table S9 and S10) were aligned by MAFFT Version 7 using the E-INS-i algorithm and BLOSUM62 scoring matrix (77). The evolutionary history was inferred using the Neighbor-Joining method (78). The evolutionary distances were computed using the Jukes Cantor genetic distance model implemented in the MEGA X software (79). A bootstrap support of ≥ 50% is given for 1000 replicates each.

## Conclusion

Zygomycetes have long been industrially used as producers of long-chain unsaturated fatty acids, but their use as resource of natural products has not been investigated yet. Here, we report on early diverging fungi as a novel resource of bioactive compounds and demonstrated that their genomes encode functional secondary metabolite genes. The two cyclopentapeptide synthetases, MpcA and MpbA, from *M. alpina* are responsible for malpicyclin and malpibaldin biosynthesis, respectively. The surprising structural and mechanistic similarity to bacterial non-ribosomal peptide synthetases (NRPS) point to an endobacterial origin of *Mortierella* NRPS genes that differ from their asco- and basidiomycete counterparts and may have evolved independently.

### Nucleotide accession number and deposit

The sequence of full-length transcripts of *mpcA* and *mpbA* were deposited at GenBank (IDs: MN800760 and MN800759).

## Acknowledgement

We are grateful to Heike Heinecke and Andrea Perner (both Hans-Knöll-Institute, Jena) for their technical assistance in recording NMR and HR-MS/MS spectra, respectively. We thank Kerstin Voigt (Jena Microbial resource Collection, Jena) for initial antimicrobial testings and Sarah Niehs (HKI Jena) for donation of genomic DNA from *R. microsporus* and *P. rhizoxinica*. Hajo Kries gratefully acknowledges a fellowship of the Daimler und Benz Foundation and financial support from the DFG and the *Fonds der Chemischen Industrie* (FCI).

